# Inhibition of Carbonic Anhydrase Using Aspirin is a Novel Method to Block Schistosomiasis Infection of the Parasitic Trematode, *Schistosoma mansoni,* in the intermediate snail host, *Biomphalaria glabrata*

**DOI:** 10.1101/2023.05.10.540221

**Authors:** Simone Parn, Gabriela Lewis, Matty Knight

**Author notes:** Corresponding author Name: Professor Matty Knight.

## Abstract

Schistosomiasis is a major public health concern worldwide. Although praziquantel is currently available as the only treatment option for schistosomiasis, the absence of reliable diagnostic and prognostic tools highlights the need for the identification and characterization of new drug targets. Recently, we identified the *B. glabrata* homolog (accession number XP_013075832.1) of human CAXIV, showing 37% amino acid sequence identity, from a BLAST search in NCBI (National Center for Biotechnology Information). Carbonic Anhydrases (CAs) are metalloenzymes that catalyze the reversible hydration/dehydration of CO_2_/HCO_3_. These enzymes are associated with many physiological processes, and their role in tumorigenesis has been widely implicated. CAs create an acidic extracellular environment that facilitates the survival, metastasis, and growth of cancer cells. In this study, we investigated the role of CA inhibition in *B. glabrata* snails exposed to *S. mansoni* miracidia. We analyzed the expression of the *B. glabrata* CA encoding transcript in juvenile susceptible and resistant snails, with and without exposure to *S. mansoni*. Our results showed that the expression of the CA mRNA encoding transcript was upregulated during early and prolonged infection in susceptible snails (BBO2), but not in the resistant BS-90 stock. Notably, sodium salicylate, a form of aspirin, inhibited the expression of CA, post-exposure, to the parasite. Increasing research between parasites and cancer has shown that schistosomes and cancer cells share similarities in their capacity to proliferate, survive, and evade host immune mechanisms. Here, we show that this model system is a potential new avenue for understanding the role of CA in the metastasis and proliferation of cancer cells. Further studies are needed to explore the potential of CA as a biomarker for infection in other schistosomiasis-causing parasites, including *S. japonicum* and *S. haematobium*.

## 1. INTRODUCTION

Schistosomiasis is a neglected tropical disease which is estimated to impact more than 600 million people in the world each year (Verjee, 2019). According to the Centers for Disease Control and Prevention, it is the second most persistent parasitic disease after malaria (Parasites, 2018). While schistosomiasis is mostly transmitted in sub-Saharan Africa, cases of its spread to Europe have been reported (Boissier et al., 2016). The parasites that cause schistosomiasis utilize the freshwater gastropod pulmonate snails as obligate hosts for the development of its asexual stages. Contact with contaminated water inadvertently leads to infection by skin penetration with infectious cercaria that are released into the water from infected snails. After penetrating the skin, cercariae lose their tail, transforming into schistosomula that develop into mature adult male and female worms that inhabit the vasculature where they mate to produce eggs that cause widespread chronic morbidity and infertility among human populations (Schistosomiasis, 2023). Specifically, infections caused by the species *Schistosoma haematobium* are known to lead to bladder cancer and female genital schistosomiasis (Hotez et al., 2019). As part of a prevention program, the World Health Organization (WHO) has proposed that school-aged children aged 5-15 years, who face the highest infection rates, receive initial parasitological assessment (WHO, 2018). Additionally, the WHO estimates that by 2030, schistosomiasis will be eliminated as a public health concern (WHO, 2020). Although several attempts have been made toward effective preventative and therapeutic measures to control the transmission of schistosomiasis, long-term reduction remains elusive. To combat the widespread disease, better diagnostic and prognostic tools, especially biomarkers to track the presence of larval parasites in the snail are of immediate necessity.

The high impact of this disease on global health burden has promoted scientific efforts to tackle the complex mechanisms underlaying successful schistosome infection. The freshwater snail *Biomphalaria glabrata,* an obligate intermediate host for the parasitic trematode *Schistosoma mansoni*, has been studied for decades at the molecular level to understand its interaction with the parasite. The genome characterization of the snail *B. glabrata* has allowed for investigation of the molecular determinants of the snail relationship with the schistosome (Adema et al., 2017). Studies of comparative immunology have enabled scientists to identify genes and pathways involved in the parasite development (Bridger et al., 2018; Raghavan et al., 2003). Additionally, differential gene expression studies have led to the identification of several cancer-and stress-related transcripts that are elevated early and significantly in the susceptible *B. glabrata* snail stocks compared to their resistant counterparts (Knight et al., 2014).

Recently, Carbonic Anhydrases (CAs) have been identified as potential targets in the development of novel drugs for parasitic diseases (Zolfaghari et al., 2022; Angeli et al., 2020). CAs are a family of zinc-containing enzymes, responsible for the reversible conversion of CO_2_ into bicarbonate and protons (Supuran et al., 2008). In humans, CAs exist in at least 15 isoforms and differ based on their enzymatic activity and cellular location: hCA I, II, III, VII, and XIII reside in the cytosol; hCA IV, IX, XII, and XIV are associated with the cell membrane; hCA VA, and VB are mitochondrial; and hCAVI is found in saliva. These enzymes are important for pH and CO_2_ homeostasis regulation, and for other biosynthesis processes, such as lipogenesis, gluconeogenesis, ureagenesis, and calcification (Pinard et al., 2015). Aberrant expression of some CA isoforms can lead to pathogenic outcomes, like carcinogenesis, obesity, and epilepsy (Poggetti et al., 2022). Notably, the expression of carbonic anhydrase IX is widely described in several hypoxic solid tumors (Svastová et al., 2004; Pastorekova et al 2019; Becker et al., 2020). In the acidic microenvironment, CAIX allows tumor cells to adapt by seizing the high CO_2_ in the form of hydrogen and bicarbonate ions.

We found through *in silico* analysis, the presence of CA in the snail *B. glabrata* genome and confirmed its evolutionary conservation. Further gene expression analysis using qPCR revealed a significant upregulation of CA in the snail upon exposure to the parasite *S. mansoni*, prompting us to further investigate its role in the snail-schistosome interaction. In this study, we investigated the expression of CA encoding transcript in susceptible and resistant *B. glabrata* stocks in response to early and prolonged exposure to *S. mansoni*. Given CA’s involvement in tumor progression and its similarity to the human enzyme, we hypothesized that the snail/schistosome relationship could mimic the activity of human CA in cancer cells. Here, we show that CA serves as a potentially useful biomarker for detecting schistosomiasis infection in infected snails, with the ability to differentiate disease from normal unaffected snails. Furthermore, we found that sodium salicylate, a water-soluble form of aspirin, inhibits the CA transcript in susceptible snail stocks. These findings provide insight into the potential role of CA in the development and progression of schistosomiasis and its potential as a therapeutic target.

## 2. MATERIALS AND METHODS

### 2.1 *Biomphalaria glabrata* Stocks

The *Biomphalaria glabrata* juvenile (3-5 mm in diameter) susceptible (BBO2) and parasite resistant (BS90) snail stocks were utilized in the study. Snails were maintained in aquaria in de-aerated water and fed with romaine lettuce twice a week.

### 2.2 Snail exposure to *S. mansoni* miracidia

Miracidia hatched from eggs isolated from 7 weeks infected mice livers were collected from the Biomedical Research Institute (Rockville, MD). Both BBO2 and BS90 juvenile snails (3mm in diameter) were exposed individually to 10-12 miracidia in 1.0 ml de-aerated water in a 6-well microtiter plate at room temperature. The snails were exposed for different time points at 0 min, 30 min, 1hr, 2hr and 4hr. Following exposure, snails were either processed in 500 µl of RNAzol immediately or kept at -80°C until needed.

### 2.3 BLAST analysis and primer design

Several bioinformatics approaches were used to test the working hypothesis. First, we searched for *B. glabrata* homologs of human related Carbonic Anhydrase encoding transcript using the protein database Uniprot (www.uniprt.org). The amino acid or FASTA file sequences were individually deposited into the Basic Local Alignment Search Tool (BLAST) (Altschul et al., 1990) to identify the *B. glabrata* homolog. The analysis was followed by a SMART BLAST analysis to validate the identity of the *B. glabrata* homolog and evolutionary relatedness to other vertebrate and invertebrate organisms’ amino acid sequence, including human. We then used the mRNA transcript of corresponding *B. glabrata* protein to design gene specific primers (Smith et al., 2021). The primers were designed excluding the *S. mansoni* corresponding CA ortholog. Oligonucleotide Primers for qPCR were obtained from Eurofins Genomics (Louisville 13 KY, 40204).

### 2.4 Trans-well *in vitro* co-culture of miracidia with *B. glabrata* embryonic cell line (BGE)

Bge cells were cultured as previously described, but in 6-well microtiter plates to confluency (Coelho et al., 2020).

### 2.5 RNA extraction

Total RNA was extracted from whole juvenile snails (unexposed and exposed) using the RNAzol method, according to the manufacturer’s instructions (Sigma-Aldrich, USA). Briefly, 500 μl of RNAzol reagent was added to the snail samples and homogenized using a pestle and a motor. Homogenate was mixed with 200 μl of sterile dH_2_O, vortexed and incubated at room temperature for 10 min before centrifugation at 13, 000 x g for 10 min at 26°C. Ethanol (75%) was added to the recovered supernatant after centrifugation and samples were incubated on ice for 10 min before another centrifugation at 13, 000 x g for 10 min at 4°C. The supernatant was removed while the pellet was left undisturbed. The pellet was washed with 70% ethanol and centrifuged at 13,000x g for 10 min at 4°C. The alcohol wash was removed, and the pellet was air-dried. RNA pellets were dissolved in 20 μl of RNase-free dH_2_O. RNA yield was determined by quantitation under UV absorbance at wavelength 260 nm using the NanoDrop 1000 spectrophotometer (Thermo Scientific). RNA samples were used either immediately or kept frozen at -80°C until needed to synthesize cDNA. Trace amounts of contaminating genomic DNA were removed by treating all RNA samples with RNAse-free DNase I prior to use for cDNA synthesis according to the manufacturer’s instructions (Promega, WI).

### 2.6 cDNA synthesis and qualitative RT-PCR

Complementary DNA was synthesized from total RNA using a cDNA synthesis kit (TermoFisher Scientific). Briefly, a master mix was prepared using 5X Reverse Transcriptase Buffer, 100 mM DTT, RNase OUT, Oligo DT, 100 mM dNTPs and Reverse transcriptase, which was aliquoted to labelled 1.5ml Eppendorf tubes. 1 ul of RNA (500ng) was used per reaction. The tubes were incubated at 42°C water bath for 2 hours and DNA concentrations were measured using NanoDrop at the wavelength of 260 nm. DNA samples were then diluted to 200ng each for qualitative PCR reaction. Experimental and reference (myoglobin) primers were diluted to 15 uM. In addition to diluted primers, the reaction was synthesized using nuclease free water, Taq Polymerase, dNTPs and Mg^+2^ 10X Buffer in a preheated thermocycler for 2 hours. Amplified PCR products were analyzed by 1.2% TBE-agarose electrophoresis.

### 2.7 Real-time quantitative RT-PCR

The real-time PCR measurement of cDNAs was performed using PowerUp SYBR Green Master Mix (Applied Biosystems) and normalized to the expression of myoglobin as standard for the housekeeping gene. The primers were: Carbonic Anhydrase forward: *caggagcagtttaggaagggc*; Carbonic Anhydrase reverse: *tcggctcaaaactcacctcc*; Myoglobin forward: *gatgttcgccaatgttccc*; Myoglobin reverse: *agcgatcaagtttccccag*. Each sample was run in technical triplicate. qRT-PCR was performed using Applied Biosystems 7300 RT PCR system, as described previously (Smith et al., 2021).

### 2.8 Treatment of susceptible snails with sodium salicylate

To determine the effect of sodium salicylate on naïve snails, snails were treated with 100ng/ml of sodium salicylate (Sigma Aldrich, St. Louis, MO) overnight (18 hours) at room temperature in 6-well microtiter plates (Smith et al., 2021). Following overnight treatment, snails were washed in nuclease free water and placed in 24-well microtiter plates. These snails were either unexposed (0 min) or individually exposed to *S. mansoni* 10-12 miracidia for 120 minutes at 25 LJC. Exposed and unexposed snails were processed for RNA as described above or kept in beakers to evaluate cercaria shedding at 4, 6 -weeks post-exposure to infection. The inhibitory concentration was investigated by means of 2-fold dilution of inhibitor.

### 2.9 Statistical analysis

Statistical analyses were performed using GraphPad Prism 8 software. All data are presented as mean ±SD. Differences between the groups were assessed using Student’s *t* test, Welch’s *t* test and 2-way analysis of variance (ANOVA) and Tukey’s test wherever relevant by comparing the differential expression (delta-Ct value) of the transcripts among treatment and control groups. Fold change was determined by utilizing uniform expression of the housekeeping myoglobin transcript as standard. Differences were considered statistically significant if *p*<0.05, with level of significance denoted as follows, ****, *p* ≤ 0.0001, ***, *p* ≤ 0.001, **, *p* ≤ 0.01, *, *p* ≤ 0.05, and ns, *p* > 0.05.

## 3. RESULTS

### 3.1 Carbonic anhydrase is highly conserved in *B. glabrata*

Based on previous RNAseq analysis (Smith et al., 2021), we identified and validated the differential expression of carbonic anhydrase protein (Accession number XP_013085564) through a BLAST analysis. Utilizing SMART BLAST analysis, we revealed the evolutionary relatedness of the Carbonic Anhydrase in *B. glabrata* to that of other organisms (**Fig. 1**). Our phylogenetics results showed that *B. glabrata* is closely related to vertebrates (house mouse, zebrafish and human) and an invertebrate fruit fly (*D. melanogaster)*, indicating co-evolution of parasitism in these organisms. Likely, the CA gene has been selected over millions of years to accommodate the parasite in these invertebrate and vertebrate hosts (**Table 1**, **Fig. 1**). Interestingly, using Clustal Omega, we aligned the CA encoding enzyme corresponding to *B. glabrata* and *S. mansoni* and observed a 34.7% amino acid identity (**Supp. Fig. 1**). Further analysis showed that the CA encoding transcript denotes a single copy gene in the *B. glabrata* genome.

**Figure 1.**
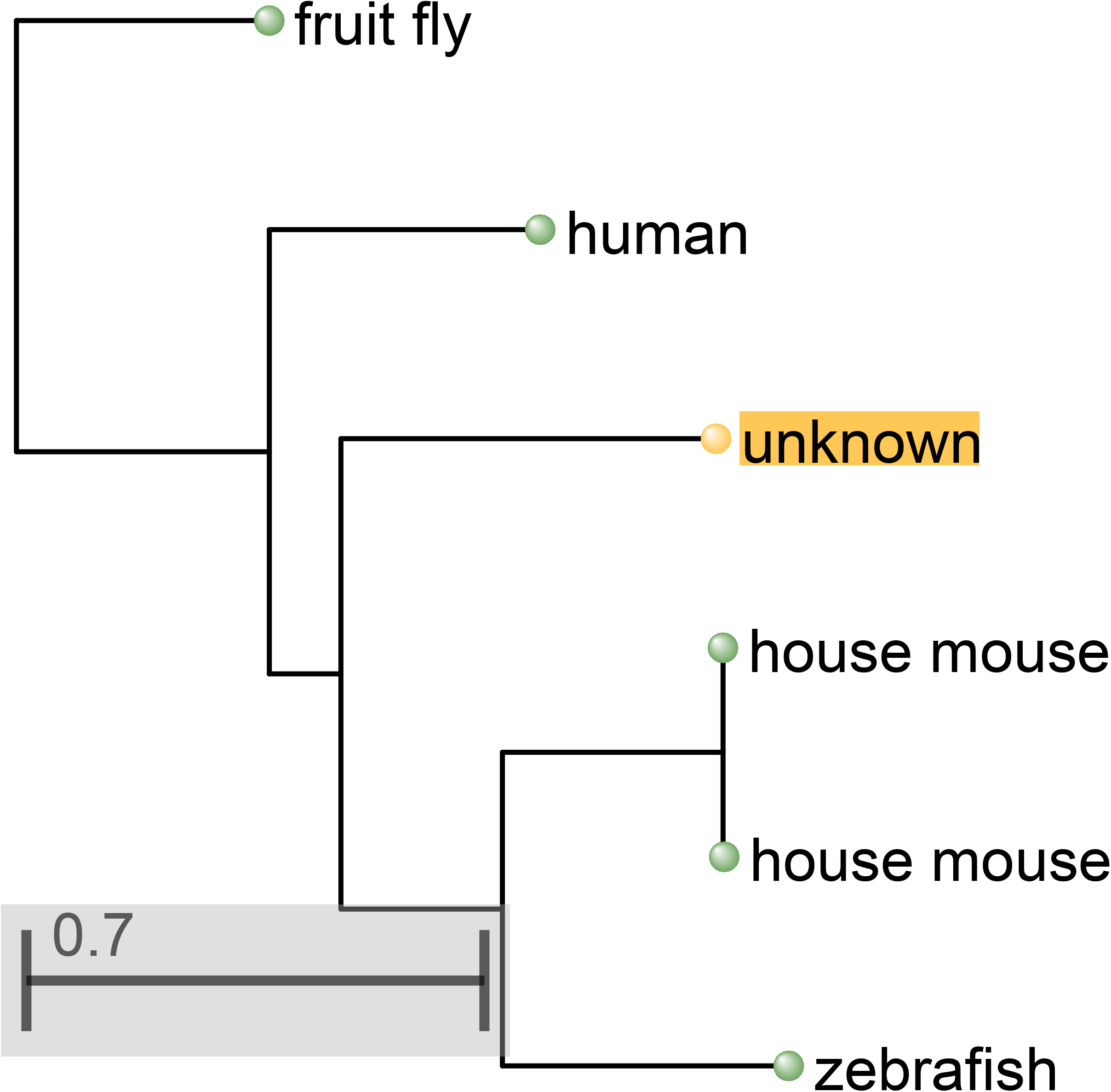
Phylogenetic tree analysis highlighting *B. glabrata* carbonic anhydrase homolog’s evolutionary relatedness to the human and other organisms. The phylogenic tree is based on amino-acid sequence alignment (ClustalW). Results indicate that *B. glabrata* is closely related to fruit fly (*D. melanogaster)* and other vertebrate organisms (house mouse, zebrafish and human). Unknown = *B. glabrata* homolog.

**Table 1.** Amino acid sequence alignment analysis of *B. glabrata* Carbonic Anhydrase 14-like protein indicates high evolutionary conservancy in relation to several organisms.

|  | Query<br>Cover | E Value | Percent<br>Identity | Accession |
| --- | --- | --- | --- | --- |
| *carbonic anhydrase 14 isoform X1*<br>[*Homo sapiens*] | 85% | 6,00E-19 | 37.82% | XP_005245116.1 |
| *carbonic anhydrase 14 precursor*<br>[*Danio rerio*] | 85% | 1,00E-17 | 34.71% | NP_001315073.1 |
| *carbonic anhydrase 1* [*Drosophila*<br>*melanogaster*] | 66% | 1,00E-17 | 45.45% | NP_523561.1 |
| *carbonic anhydrase 12 isoform X1*<br>[*Mus musculus*] | 71% | 3,00E-17 | 41.18% | XP_006511611.1 |
| *carbonic anhydrase 14 isoform X6*<br>[*Homo sapiens*] | 71% | 4,00E-17 | 43.00% | XP_016856382.1 |
| *carbonic anhydrase* [*Schistosoma*<br>*mansoni*] | 86% | 3,00E-31 | 34.70% | QDD67328 |

### 3.2 Carbonic anhydrase is a surrogate marker of infection

To determine whether changes in the expression of CA occur in juvenile *B. glabrata* snails after parasite exposure, qPCR was utilized to measure temporal changes at different time-points after exposure to *S. mansoni* miracidia. Results showed that CA was differentially expressed between susceptible (BBO2) and resistant (BS-90) snail stocks upon early exposure to the parasite. Thus, a significant upregulation (1.42-fold) was observed following 2h post-exposure, increasing to 1.77-fold after 4hr infection in the susceptible snail (**Fig. 2**). Moreover, sustained expression of CA mRNA was observed throughout the prolonged 6-week infection period of the susceptible snail (**Fig. 3**). The CA mRNA transcript was significantly upregulated between 1 to 6 weeks (1.9-to 8-fold) post-exposure in the susceptible BBO2 snails but not in the resistant BS90 snails. To further test the expression of CA, we validated the results using the *B. glabrata* embryonic cell line (BGE) co-cultured with *S. mansoni* miracidia. Our results showed that CA was upregulated by 1.1-fold at 0.5-hour and by 1.4-fold following 2-hour exposure to miracidia (**Fig. 4**). Interestingly, similar to CA expression in susceptible BBO2 snails, CA expression in BGE cell line alone was down-regulated at the 1-hour mark.

**Figure 2.**
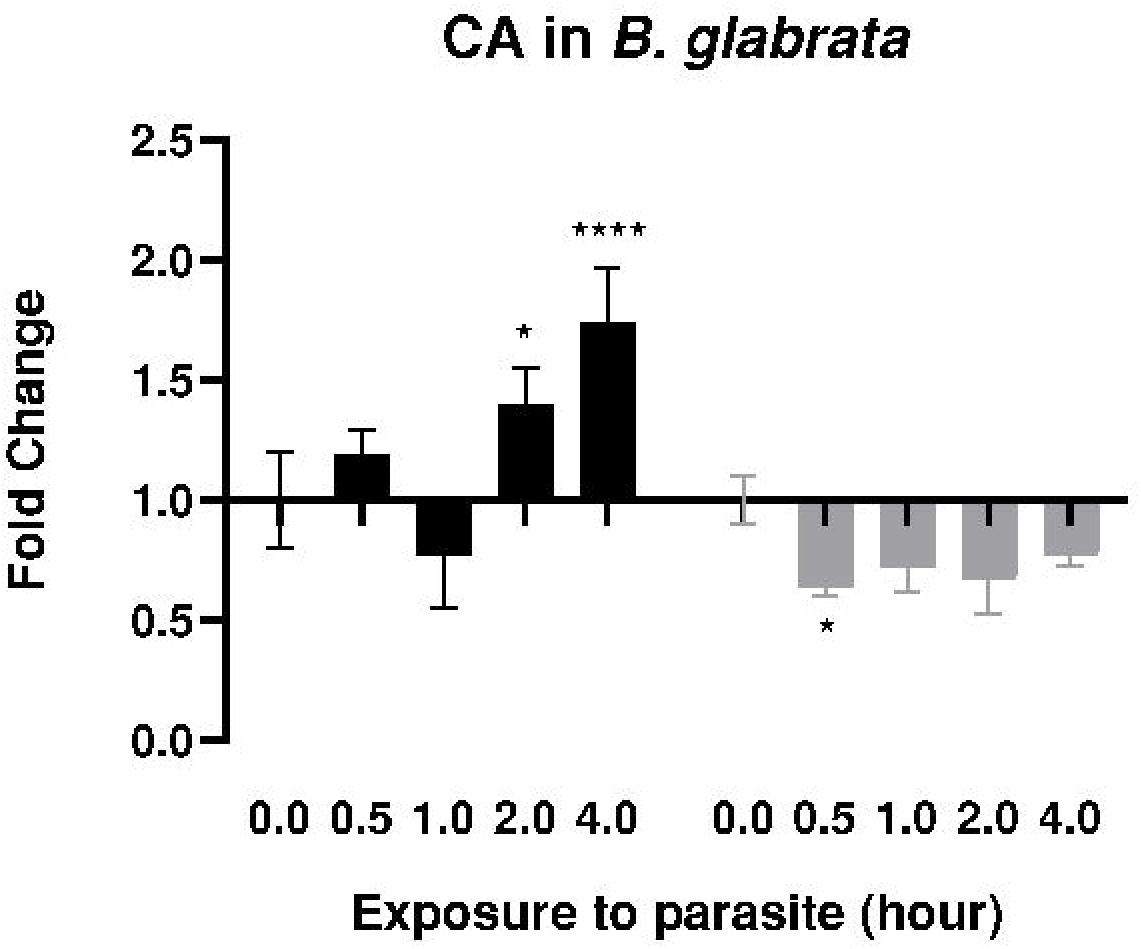
qPCR analysis of RNA from susceptible BBO2 (black histograms) and resistant BS-90 (gray histograms) juvenile snails exposed for 30 min, 1 hour, 2 hours and 4 hours to the parasite *S. mansoni* miracidia. Histograms show the expression of CA encoding transcript in *B. glabrata* snails at specific time points from four biological replicates (the total snails used for four biological replicates N=40). Note the increase in fold change in susceptible BBO2 snails compared to the resistant BS90 snails after parasite exposure. Fold change was determined as described previously by utilizing uniform expression of the reference transcript, ****, p ≤ 0.0001, ***, p ≤ 0.001, **, p ≤ 0.01, *, p ≤ 0.05, and ns, p > 0.05.

**Figure 3.**
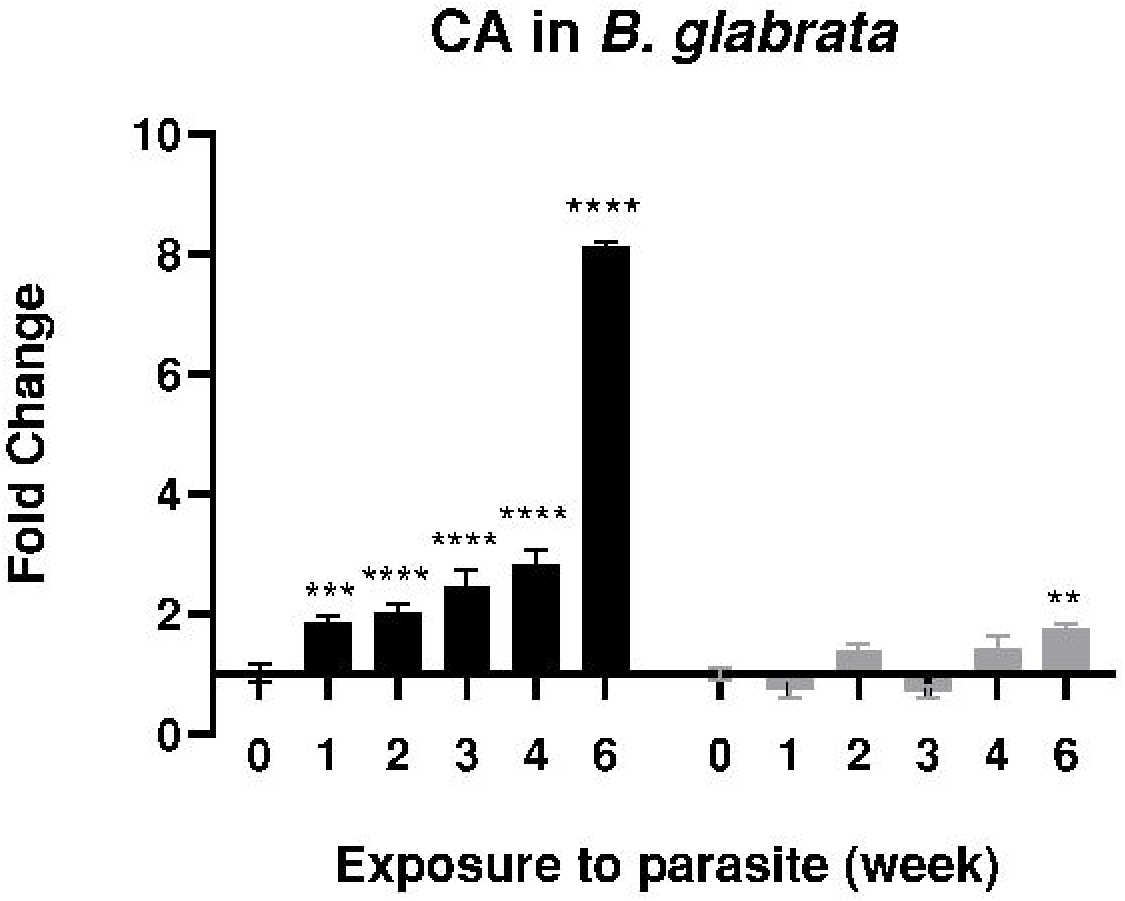
qPCR analysis of RNA from susceptible BBO2 (black histograms) and resistant BS-90 (gray histograms) juvenile snails exposed for 2 hours and kept up to 6 weeks to the parasite *S. mansoni*. Histograms show expression of CA transcripts in juvenile snails from three biological replicates (N=36). Non-exposed (0min) juvenile snails served as controls. A significant increase in fold change can be observed in the susceptible BBO2 snails compared to the resistant BS90 snails. ****, p ≤ 0.0001, ***, p ≤ 0.001, **, p ≤ 0.01, *, p ≤ 0.05, and ns, p > 0.05.

**Figure 4.**
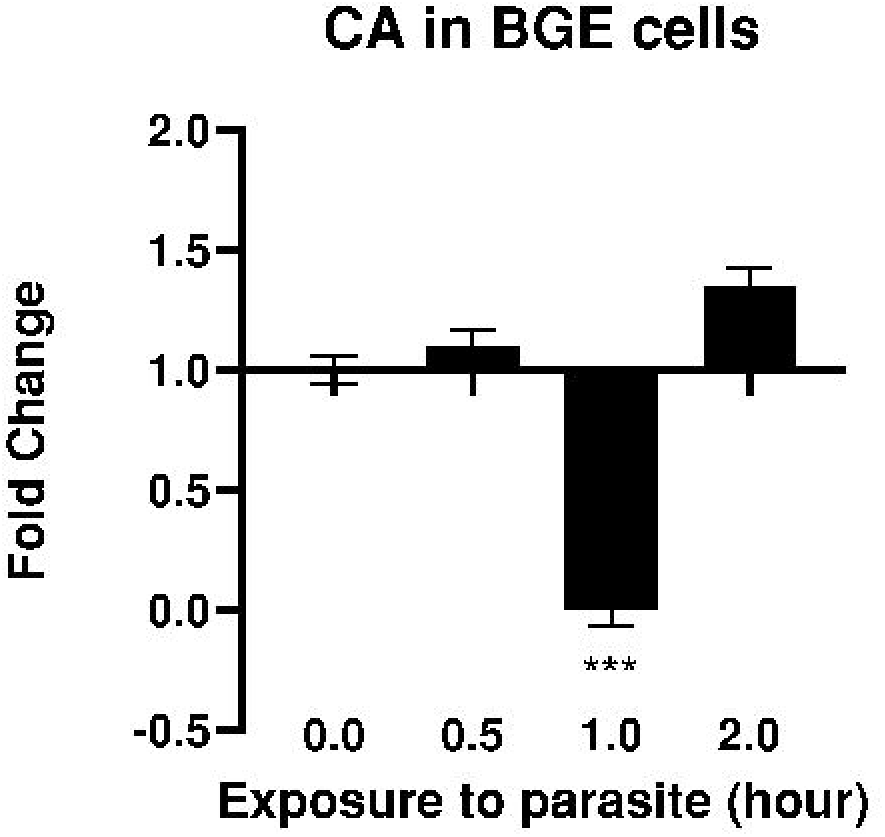
qPCR analysis of RNA from *B. glabrata* embryonic cell line (BGE) co-cultured with *S. mansoni* miracidia. BGE cells were exposed to *S. mansoni* miracidia for 30 min, 1 hour and 2 hours (N=8). Non-exposed (0min) BGE cells served as controls. ****, p ≤ 0.0001, ***, p ≤ 0.001, **, p ≤ 0.01, *, p ≤ 0.05, and ns, p > 0.05.

### 3.3 Sodium salicylate is a potent inhibitor of carbonic anhydrase

We also investigated the potential of aspirin to inhibit CA expression in susceptible BBO2 snails. To achieve this, we utilized sodium salicylate, a major metabolite of aspirin, which is a water soluble non-steroidal anti-inflammatory agent known to induce apoptosis of cancer cells and reduce tumor growth (National Center for Biotechnology Information, 2023). Treatment of BBO2 snails with 100 ng/ml sodium salicylate overnight (18 hours) prior to schistosome exposure was shown to suppress CA expression (**Fig. 5**). Through a 2-fold serial dilution assay of the drug, we determined the required inhibitory concentration of sodium salicylate for suppressing CA expression. Our findings, as seen from **Figure 6**, consistently showed downregulation of CA encoding RNA transcript. Notably, a concentration as low as 12.5 ng/mL of sodium salicylate was sufficient to inhibit CA. Importantly, CA expression remained downregulated in sodium salicylate treated snails with exposure to the parasite *S. mansoni*.

**Figure 5.**
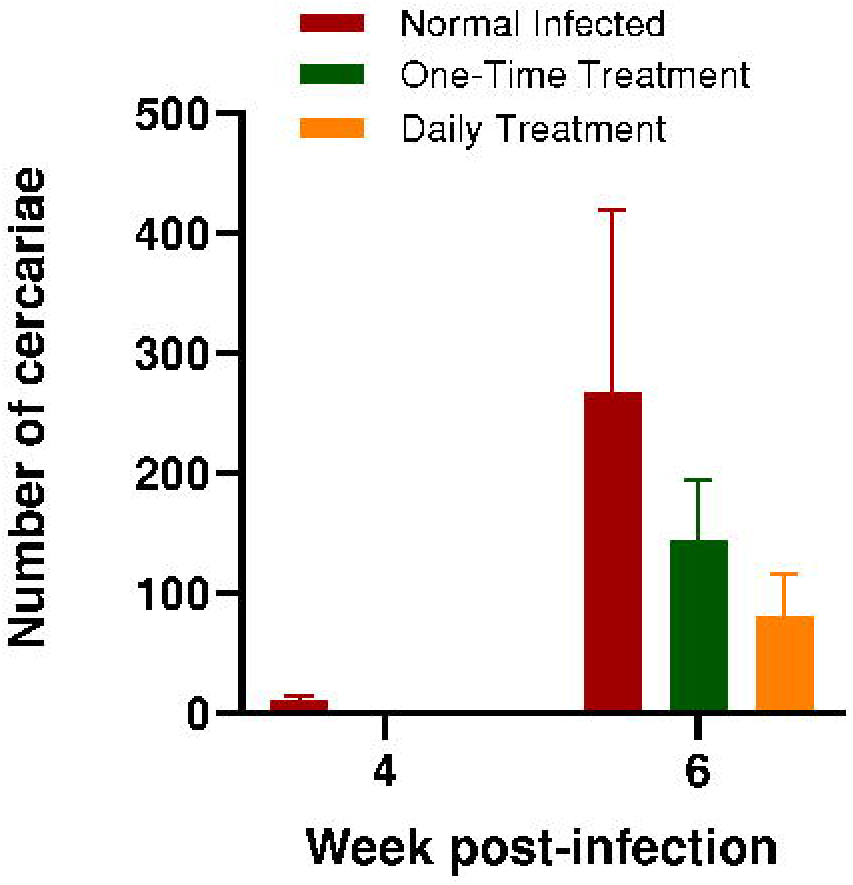
Schistosome recovery from infected *B. glabrata* following treatment with 100 ng/ml sodium salicylate. Susceptible BBO2 snails were treated overnight with 100 ng/mL sodium salicylate and exposed to *S. mansoni* miracidia (10-12 miracidia per snail) for two hours. The histograms show cercarial shedding in non-drug treated/infected snails (red); one-time drug treated (16 hours)/infected snails (green); and daily drug-treated/infected snails (yellow) from four biological replicates (N=12). The fold changes of the histograms were noted as non-significant (ns) through 2-way ANOVA analysis.

**Figure 6.**
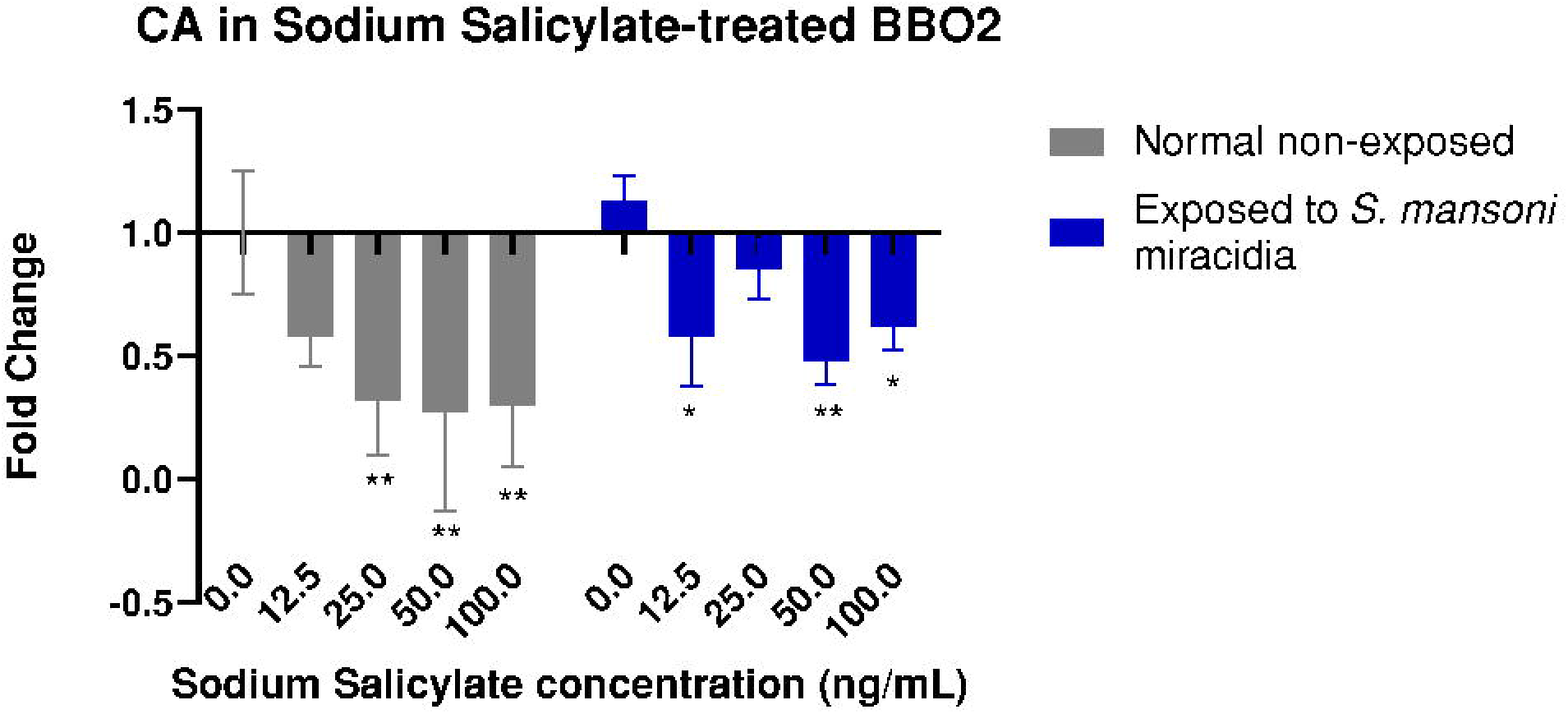
qPCR analysis of RNA from susceptible juvenile BBO2 snails unexposed (gray histograms) and exposed (blue histograms) for 2 to the parasite *S. mansoni* miracidia. Histograms show expression of carbonic anhydrase encoding transcript in normal non-drug treated juvenile snails (0 ng/ml) and those treated with sodium salicylate (12.5 ng/ml, 25 ng/ml, 50 ng/ml or 100 ng/ml) from three biological replicates (N=30). Non-exposed (0min) and non-drug treated snails served as controls. Note the upregulation of CA transcript in non-drug treated exposed snails and the downregulation in non-exposed and exposed drug-treated snail groups. ****, p ≤ 0.0001, ***, p ≤ 0.001, **, p ≤ 0.01, *, p ≤ 0.05, and ns, p > 0.05.

### 3.4 Sodium salicylate may reduce infection burden in susceptible *B. glabrata* snails

To test the effects of suppression of CA gene *in vivo*, sodium salicylate treated BBO2 snails were maintained after exposure to parasite to examine for cercariae shedding after 4-6 weeks. Our results indicated that a one-time overnight treatment with 100 ng/mL sodium salicylate prior to schistosome infection (2-hour exposure) was insufficient to block infection in the snails. While neither the non-treated infected snails nor the drug-treated infected snails shed at 4 weeks, following 6 weeks’ time, all snails shed cercaria (**Fig. 5**). Non drug-treated infected control snails shed the highest number of cercariae. Similarly, one-time drug treated snails shed cercariae, although at lower levels. Following the experiment, we repeated the sodium salicylate treatment in susceptible BBO2 snails, this time maintaining the infected snails continuously in 100 ng/ml sodium salicylate both before (overnight) and after 2-hour exposure to the parasite. Following 4 weeks of exposure, we observed a decrease (1 cercariae shed) or complete absence of cercarial shedding in the treated/exposed snails compared to the non-treated/exposed control group which shed an average of 9 cercaria. Surprisingly, cercarial shedding was observed in the daily drug-treated infected snails at 6 weeks, however, at lower levels compared to the non-drug treated infected control group and one-time drug-treated infected snails (**Fig. 5**).

## 4. DISCUSSION

Schistosomiasis remains a serious public health concern, especially in many low-income sub-Saharan African areas. To date, however, the only known effective treatment against the causative agent of schistosomiasis-the parasitic trematode, is praziquantel (Schistosomiasis, 2023). Schistosomiasis is endemic in 78 countries and at least 600 million people are at risk of becoming infected every year (Schistosomiasis, 2023). The WHO has announced that by 2030 schistosomiasis will be eradicated (WHO, 2020). However, without a preventative vaccine, and only a single therapeutic drug currently available, this might prove elusive. Efforts to control schistosomiasis have been attempted by a mass drug chemotherapy program that provided treatment for school aged children (Kokaliaris et al., 2022). The detection of infection in patients relies on the detection of parasite eggs in stool (*S. mansoni*, and *S. japonicum*) and urine (*S. hematobium*) (Schistosomiasis, 2023). For the past 18 years this method of controlling schistosomiasis, although successful, has not eliminated the disease. This is because the parasite is difficult to completely eliminate without attention to the transmission in the intermediate snail host.

The control of the freshwater snail has been made possible largely by the use of molluscicides that can be environmentally toxic as well as disturbing fragile ecosystems. There are currently no reliable convenient biomarkers available for the detection of the parasite in infected snails. Previous studies have described the cloning and characterization of 121-bp tandem DNA repeat sequences in the schistosomal genome (Hamburger et al., 1987). Although, the application of such DNA based probes presents a high detection sensitivity, the degree of specificity has not been addressed. Furthermore, these studies have not demonstrated the specific time-points at which the infection in snails can be detected. Results from our study show that the presence of carbonic anhydrase is a good and accurate biomarker for early and prolonged detection of *S. mansoni* in the intermediate snail host*, B. glabrata*.

From qPCR analysis, temporal regulation during the initial 30 minutes to 4 hours post exposure of the snail to the parasite revealed that CA is upregulated in susceptible BBO2 snails during early infection but downregulated in their resistant counterparts. Given that CAs are predominantly involved in hypoxia and pH regulation in cancer cells, we observed a significant upregulation of CA expression in susceptible snail lines, with a 1.42-fold increase after two hours of exposure and 1.77-fold increase post four hours (**Fig. 2**). Moreover, we found that CA mRNA levels were further increased in susceptible snails during prolonged infection up to 6 weeks (**Fig. 3**), indicating the critical role of CA in upholding the proliferative function. To further validate the results, we used the *B. glabrata* embryonic cell line (BGE) co-cultured with *S. mansoni* miracidia and observed a 1.4-fold upregulation of CA transcripts at the two-hour infection period (**Fig. 4**). Our previous research has demonstrated that transcripts encoding the heat shock proteins Hsp70 and Hsp90, including the reverse transcriptase (RT) domain, the non-LTR-transposon nimbus, are similarly upregulated in the susceptible snails but not in resistant (Ittiprasert et al., 2009). Further studies from spatial repositioning of gene loci have confirmed that schistosomes orchestrate transcription of Hsp70 transcript in susceptible *B. glabrata* snails quickly after infection (Arican-Goktas et al., 2014).

CAs also serve as important therapeutic targets in various biological processes, including acid-base balance, inflammation and angiogenesis. In humans, there are 15 CA isoforms (alpha-class CAs), with CA IX and CA XII having been linked to cancer (Supuran et al., 2008). These enzymes are transmembrane isoforms with an extracellular catalytic domain, showing high expression in solid tumors and low expression in normal tissues. The overexpression of CA IX and CA XII in tumors is associated with the survival and proliferation of cancer cells, making them attractive targets for cancer therapy (Mahon et al., 2015). In this study, we specifically examined CA isoform XIV, which has a medium-low catalytic activity similar to that of hCA XII (Nishimori et al., 2005). This isoform is highly abundant in the brain, kidneys, colon, small intestine, urinary bladder, liver, and spinal cord (Alterio et al., 2014). A Clustal Omega sequence alignment (Sievers et al., 2011) showed high amino acid conservation among the human α-CAs, specifically in the catalytic sites. Results indicated that the active site of these enzymes are highly superimposable, suggesting similar evolutionary conserved enzymatic activity among the CA encoding transcripts between the snail homolog and the other organisms. Of particular interest, structural comparison of the hCA IX catalytic domain with the CA 14-like *B. glabrata* homolog yielded a substantial sequence homology (**Supp. Fig. 2**). While the two enzyme orthologs share 35.97% sequence identity in the catalytic sites, further interrogations of the sequences revealed that the two proteins consist of identical zinc binding sites (ion binding), specifically, the three conserved histidine residues (His94, His96, and His119). This homology suggests that the isoforms may share similar functions, and one isoform may be able to compensate for the loss of the other (Aggarwal et al., 2013).

We have shown there is a link between metastatic cancers and parasitic diseases, highlighting the snail host/parasite relationship as a valuable animal model for studying the regulation of cancer-associated transcripts. Similar to cancer, the schistosome survives in the blood stream and evades the host immune system mechanisms. Deposition of *S. mansoni* eggs also causes acute and chronic inflammation of the colorectal mucosa (McManus et al., 2018). Our results indicate that CA plays a key role in the progression of schistosomiasis infection in the snail, with prolonged infection leading to increased expression of CA transcripts. CA’s role in facilitating the transport of CO_2_ across the cell membrane creating acidic environment, is one of the hallmarks of cancer. In accord with studies from CA in cancer, several human CA isoforms have been shown to be increased in tumor tissue, correlating with cancer growth and survival (Ning et al., 2022; McDonald et al., 2019; Hsieh et al., 2022; Schmidt et al., 2021). This supports the notion that the snail-schistosome model can provide useful insights into the mechanisms underlaying both parasitic infections and cancer (manuscript in preparation).

Targeting CA has emerged as a promising therapeutic strategy for cancer treatment with numerous CA inhibitors currently undergoing preclinical and clinical trials (Mussi et al., 2022; Dvořanová et al., 2020). It has been well established that a positive correlation exists between risk of cancer and aspirin intake. As part of this, several clinical trials have investigated the use of aspirin as an adjuvant therapy for cancer patients, particularly those with colorectal cancer (Sostres et al., 2014; Rothwell et al., 2010). We show in this study that sodium salicylate, a major subunit of aspirin, can inhibit CA in susceptible BBO2 snails. Although our results from qPCR show that sodium salicylate down regulates the transcript encoding carbonic anhydrase, we found that a single dose of 100 ng/ml sodium salicylate is not sufficient to block infection, likely due to its metabolism and short half-life. However, daily administration of 100 ng/ml sodium salicylate can further reduce the infection burden compared to a single dose and normal infected snails. Combination therapy involving aspirin may offer greater therapeutic benefits, as evidenced by Feitosa, et al. demonstrating a reduction in parasite load in *S. mansoni* mice (Feitosa et al., 2018). Furthermore, the combination of paraziquantel and aspirin has been shown to decrease liver pathology, highlighting the potential of this treatment approach (Sudsarn et al., 2016).

Our present work demonstrates the feasibility of detecting early and prolonged *S. mansoni* infection in snails by employing a simple and accurate procedure using qPCR. However, our study has some limitations that need to be taken into account before considering the application of CA for field studies. Firstly, considering the role of CA in acid-base regulation and creating acidic environments, change in CA activity and pH upon infection remains to be investigated further with more experiments. Secondly, to gain a better understanding of the mechanism of action of CA, further studies to determine its enzymatic activity are needed. Thirdly, the degree of enzyme activity remains to be evaluated in other schistosomiasis-causing trematodes, *S. japonicum* and *S. haematobium.* In addition, antibody detection assays are being developed to make the use of CA for rapid and practical use in the field to detect occurrence of infected snails in endemic high prevalence area before using molluscicides which can be expensive to use on a wide scale.

In summary, we have shown in this study that CA overexpression is correlated with infection and parasite progression in the snail host. We provide the first evidence of CA in mollusk species upon infection with *S. mansoni* and our research shows that treatment with salicylic acid is a competent method of inhibiting CA activity in the *B. glabrata* snail, thus inhibiting the infection of the snail with the *S. mansoni* parasite. These results indicate that CA is a promising biomarker to track schistosomiasis in early and prolonged infection stages.

## Acknowledgements

We thank Dr. Margaret Mentink-Kane at the NIAID Schistosomiasis Resource Center of the Biomedical Research Institute for the snail and parasite material used in this study. We also thank Dr. Carolyn Cousin for resources and support. This work was funded by the Clement BT Knight Cancer Foundation.

## CRediT authorship contribution statement

**Simone Parn:** Investigation, Methodology, Data Curation, Formal Analysis, Validation, Visualization, Writing – Original Draft, Writing – Review & Editing; **Gabriela Lewis:** Investigation, Methodology, Data Curation, Validation, Writing – Review & Editing; **Matty Knight:** Conceptualization, Methodology, Investigation, Resources, Project Administration, Supervision, Writing – Original Draft, Writing – Review & Editing.

## Data Availability

Data will be made available on request.

---

**Supplementary Figure 1.**
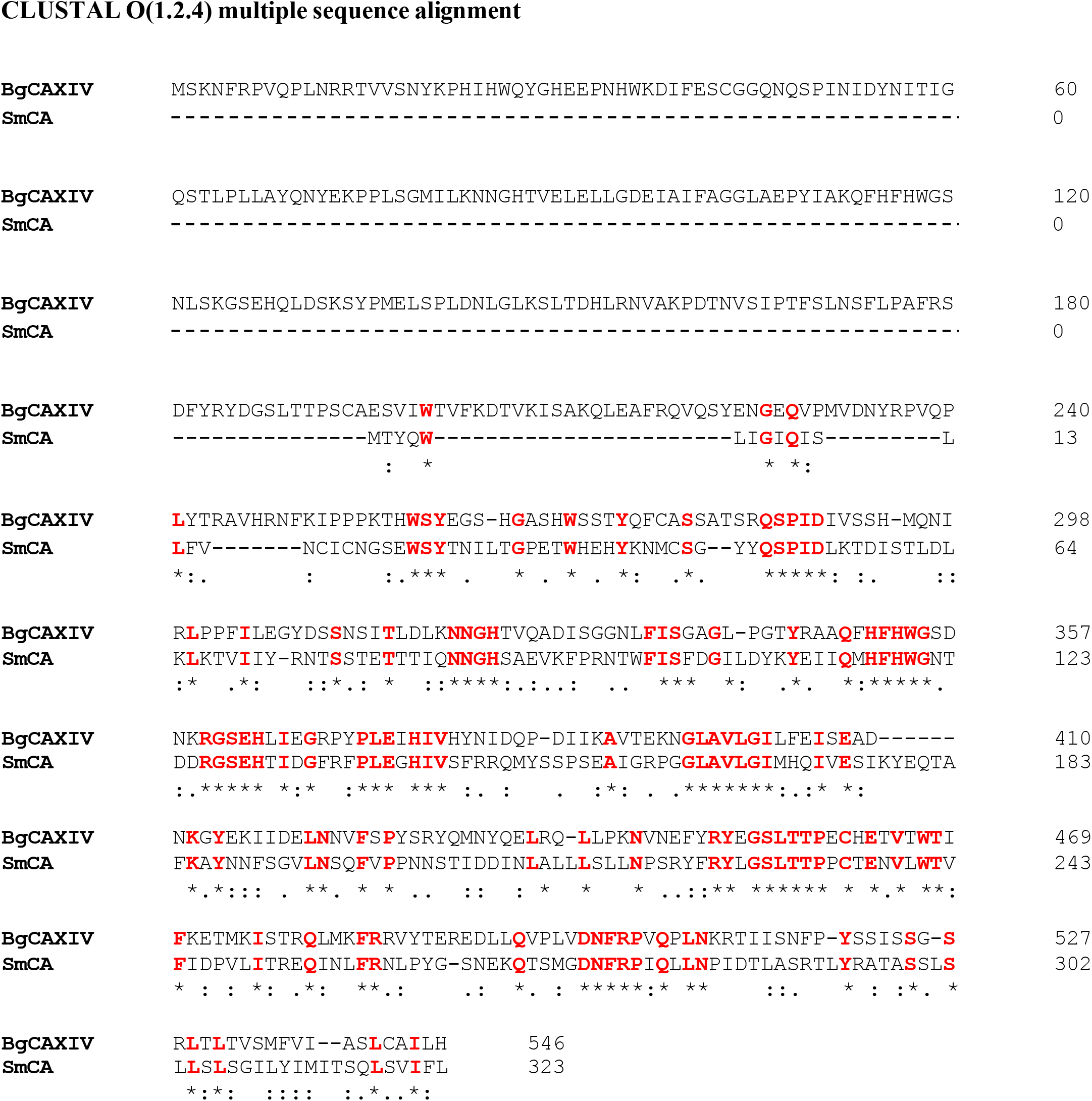
Multiple sequence alignment of the carbonic anhydrase enzyme between *B. glabrata* (accession number XP_013085564.1) and *S. mansoni* (accession number QDD67328.1) obtained from Clustal Omega. CA shares a 34.7% identity between the snail and the parasite *S. mansoni*. 100% amino acid match is indicated by an asterisk (letters marked in red).

**Supplementary Figure 2.**
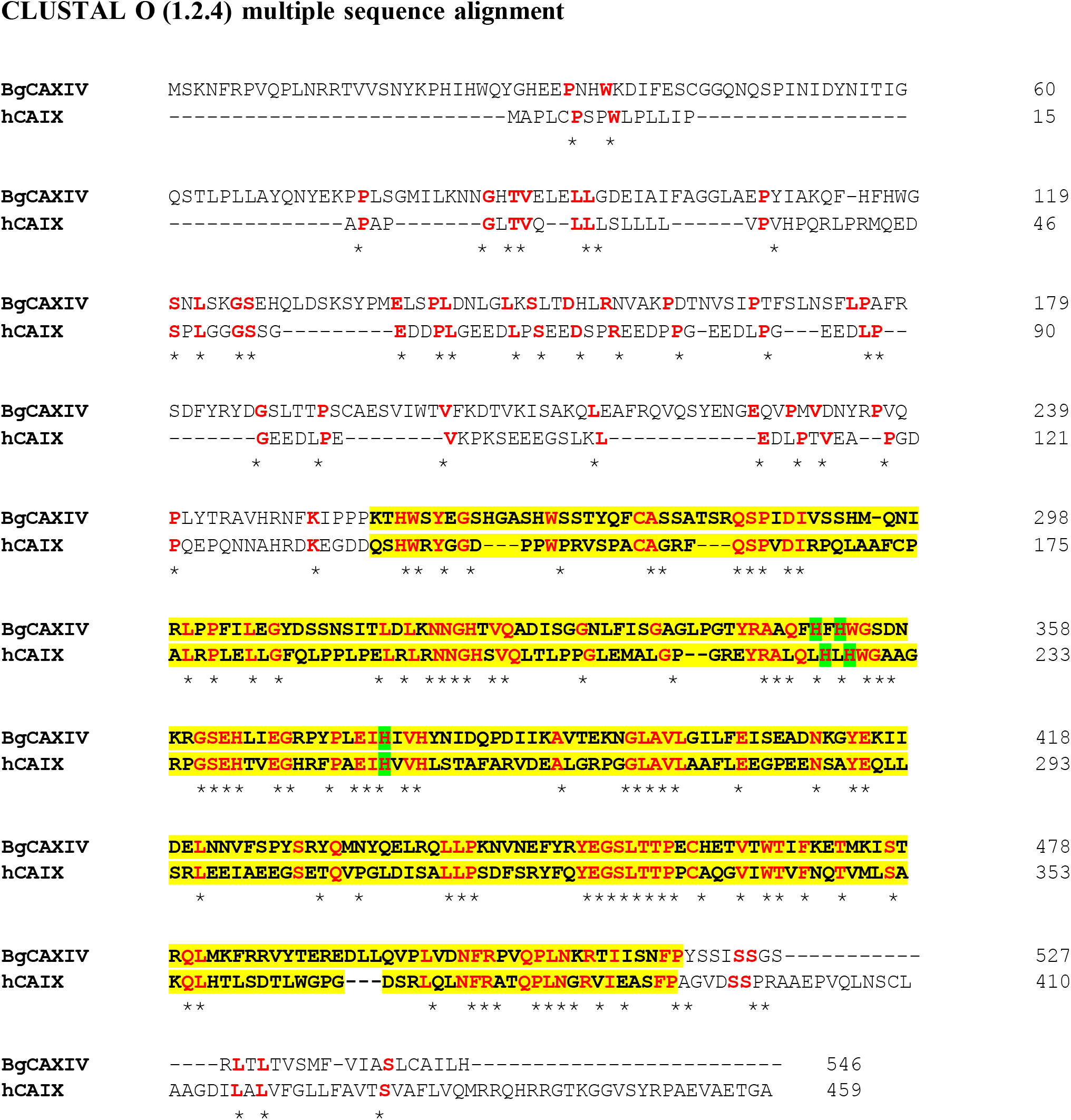
Structure-based amino acid sequence alignment of the human Carbonic Anhydrase IX (accession number: Q16790) and *B. glabrata* Carbonic Anhydrase 14-like protein homolog (accession number XP_013085564.1) obtained from Clustal Omega. The two enzyme orthologs share 31.25% sequence identity and 35.97% identity in the catalytic sites (highlighted in yellow). 100% amino acid match is indicated by an asterisk (letters marked in red). Zinc-ion binding sites are shown in green.

## REFERENCES

1. Adema C, Hillier L, Jones C. et al. Whole genome analysis of a schistosomiasis-transmitting freshwater snail. Nat Commun 8. 2017; 15451. 10.1038/ncomms15451. Accessed February 25, 2023.

2. Aggarwal M, Boone CD, Kondeti B, McKenna R. Structural annotation of human carbonic anhydrases. Journal of Enzyme Inhibition and Medicinal Chemistry. 2013, 28:2, 267–277, Doi: 10.3109/14756366.2012.737323. Accessed April 10, 2023.

3. Alterio V, Pan P, Parkkila S, Buonanno M, Supuran CT, Monti SM, De Simone G. The structural comparison between membrane-associated human carbonic anhydrases provides insights into drug design of selective inhibitors. Biopolymers. 2014;101(7):769–78. doi: 10.1002/bip.22456. Accessed April 4, 2023.

4. Altschul SF, Gish W, Miller W, Myers EW, Lipman DJ. Basic local alignment search tool. J Mol Biol. 1990;215(3):403–10. doi: 10.1016/S0022-2836(05)80360-2. Accessed April 13, 2023.

5. Angeli A, Pinteala M, Maier SS, Simionescu BC, Da’dara AA, Skelly PJ, Supuran CT. Sulfonamide Inhibition Studies of an α-Carbonic Anhydrase from *Schistosoma mansoni*, a Platyhelminth Parasite Responsible for Schistosomiasis. Int J Mol Sci. 2020;21(5):1842. doi: 10.3390/ijms21051842. Accessed April 11, 2023.

6. Arican-Goktas HD, Ittiprasert W, Bridger JM, Knight M. Differential spatial repositioning of activated genes in Biomphalaria glabrata snails infected with Schistosoma mansoni. PLoS Negl Trop Dis. 2014;8(9):e3013. doi: 10.1371/journal.pntd.0003013. Accessed March 30, 2023.

7. Becker HM. Carbonic anhydrase IX and acid transport in cancer. Br J Cancer. 2020; 122, 157–167. 10.1038/s41416-019-0642-z. Accessed April 13, 2023.

8. Boissier J, Grech-Angelini S, Webster BL, Allienne JF, Huyse T, Mas-Coma S, Toulza E, Barré-Cardi H, Rollinson D, Kincaid-Smith J, Oleaga A, Galinier R, Foata J, Rognon A, Berry A, Mouahid G, Henneron R, Moné H, Noel H, Mitta G. Outbreak of urogenital schistosomiasis in Corsica (France): an epidemiological case study. Lancet Infect Dis. 2016 Aug;16(8):971–9. doi: 10.1016/S1473-3099(16)00175-4. Accessed April 8, 2023.

9. Bridger JM, Brindley PJ, Knight M. The snail *Biomphalaria glabrata* as a model to interrogate the molecular basis of complex human diseases. PLoS Negl Trop Dis. 2018,12(8): e0006552. 10.1371/journal.pntd.0006552. Accessed April 12, 2023.

10. Coelho, F.S., Rodpai, R., Miller, A. et al. Diminished adherence of *Biomphalaria glabrata*embryonic cell line to sporocysts of *Schistosoma mansoni* following programmed knockout of the allograft inflammatory factor. Parasites Vectors 13, 511 (2020). 10.1186/s13071-020-04384-9.

11. Dvořanová J, Kugler M, Holub J, Šícha V, Das V, Nekvinda J, El Anwar S, Havránek M, Pospíšilová K, Fábry M, Král V, Medvedíková M, Matějková S, Lišková B, Gurská S, Džubák P, Brynda J, Hajdúch M, Grüner B, Řezáčová P. Sulfonamido carboranes as highly selective inhibitors of cancer-specific carbonic anhydrase IX. Eur J Med Chem. 2020; 200:112460. doi: 10.1016/j.ejmech.2020.112460. Accessed April 3, 2023.

12. Ending the neglect to attain the Sustainable Development Goals: A road map for neglected tropical diseases 2021–2030. Geneva: World Health Organization; 2020. Licence: CC BY-NC-SA 3.0 IGO. Accessed April 10, 2023.

13. Feitosa KA, Zaia MG, Rodrigues V, Castro CA, Correia RO, Pinto FG, Rossi KNZP, Avó LRS, Afonso A, Anibal FF. *Menthol and Menthone* Associated with Acetylsalicylic Acid and Their Relation to the Hepatic Fibrosis in *Schistosoma mansoni* Infected Mice. Front Pharmacol. 2018 Jan 18;8:1000. doi: 10.3389/fphar.2017.01000. Accessed April 14, 2023.

14. Hamburger J, Weil M & Pollack Y. Detection of Schistosoma mansoni DNA in extracts of whole individual snails by dot hybridization. Parasitol Res, 1987, 74, 97–100- doi: 10.1007/BF00534940.

15. Hotez PJ, Engels D, Gyapong M, Ducker C, Malecela MN. Female Genital Schistosomiasis. N Engl J Med. 2019 Dec 26;381(26):2493-2495. doi: 10.1056/NEJMp1914709. Accessed March 30, 2023.

16. Hsieh M, Huang PJ, Chou PY, Wang SW, Lu HC, Su WW, Chung YC, Wu MH. Carbonic Anhydrase VIII (CAVIII) Gene Mediated Colorectal Cancer Growth and Angiogenesis through Mediated miRNA 16-5p. Biomedicines. 2022 Apr 29;10(5):1030. doi: 10.3390/biomedicines10051030. Accessed April 3, 2023.

17. Ittiprasert W, Nene R, Miller A, Raghavan N, Lewis F, Hodgson J, Knight M. Schistosoma mansoni infection of juvenile Biomphalaria glabrata induces a differential stress response between resistant and susceptible snails. Exp Parasitol. 2009 Nov;123(3):203–11. doi: 10.1016/j.exppara.2009.07.015. Accessed March 30, 2023.

18. Knight M, Arican-Goktas HD, Ittiprasert W, Odoemelam EC, Miller AN, Bridger JM. Schistosomes and snails: a molecular encounter. Front Genet. 2014 Jul 21;5:230. doi: 10.3389/fgene.2014.00230. Accessed April 10, 2023.

19. Kokaliaris C, Garba A, Matuska M, Bronzan RN, Colley DG, et al. Effect of preventive chemotherapy with praziquantel on schistosomiasis among school-aged children in sub-Saharan Africa: a spatiotemporal modelling study. Lancet Infect Dis. 2022 Jan;22(1):136–149. doi: 10.1016/S1473-3099(21)00090-6. Epub 2021 Dec 2. Erratum in: Lancet Infect Dis. 2022 Jan;22(1):e1. Accessed April 13, 2023.

20. Mahon BP, Pinard MA, McKenna R. Targeting carbonic anhydrase IX activity and expression. Molecules. 2015 Jan 30;20(2):2323–48. doi: 10.3390/molecules20022323. Accessed April 13, 2023.

21. McDonald PC, Chafe SC, Brown WS, Saberi S, Swayampakula M, Venkateswaran G, Nemirovsky O, Gillespie JA, Karasinska JM, Kalloger SE, Supuran CT, Schaeffer DF, Bashashati A, Shah SP, Topham JT, Yapp DT, Li J, Renouf DJ, Stanger BZ, Dedhar S. Regulation of pH by Carbonic Anhydrase 9 Mediates Survival of Pancreatic Cancer Cells With Activated KRAS in Response to Hypoxia. Gastroenterology. 2019 Sep;157(3):823–837. doi: 10.1053/j.gastro.2019.05.004. Accessed April 3, 2023.

22. McManus, D.P., Dunne, D.W., Sacko, M. et al. Schistosomiasis. Nat Rev Dis Primers 4, 13 (2018). 10.1038/s41572-018-0013-8. Accessed April 13, 2023.

23. Mussi S, Rezzola S, Chiodelli P, Nocentini A, Supuran CT, Ronca R. Antiproliferative effects of sulphonamide carbonic anhydrase inhibitors C18, SLC-0111 and acetazolamide on bladder, glioblastoma and pancreatic cancer cell lines. J Enzyme Inhib Med Chem. 2022 Dec;37(1):280–286. doi: 10.1080/14756366.2021.2004592. Accessed April 3, 2023.

24. National Center for Biotechnology Information (2023). PubChem Compound Summary for CID 16760658, Sodium Salicylate. Accessed April 13, 2023.

25. Ning WR, Jiang D, Liu XC, Huang YF, Peng ZP, Jiang ZZ, Kang T, Zhuang SM, Wu Y, Zheng L. Carbonic anhydrase XII mediates the survival and prometastatic functions of macrophages in human hepatocellular carcinoma. J Clin Invest. 2022 Apr 1;132(7):e153110. doi: 10.1172/JCI153110. Accessed April 3, 2023.

26. Nishimori I, Vullo D, Innocenti A, Scozzafava A, Mastrolorenzo A, Supuran CT. Carbonic anhydrase inhibitors: inhibition of the transmembrane isozyme XIV with sulfonamides. Bioorg Med Chem Lett. 2005 Sep 1;15(17):3828–33. doi: 10.1016/j.bmcl.2005.06.055. Accessed April 10, 2023.

27. Parasites-schistosomiasis. Centers for Disease Control and Prevention. 2018. https://www.cdc.gov/parasites/schistosomiasis/index.html. Accessed April 8, 2023.

28. Pastorekova S, Gillies RJ. The role of carbonic anhydrase IX in cancer development: links to hypoxia, acidosis, and beyond. Cancer Metastasis Rev. 2019 Jun;38(1-2):65–77. doi: 10.1007/s10555-019-09799-0. Accessed March 27, 2023.

29. Pinard MA, Mahon B, McKenna R. Probing the surface of human carbonic anhydrase for clues towards the design of isoform specific inhibitors. Biomed Res Int. 2015;2015:453543. doi: 10.1155/2015/453543. Accessed April 13, 2023.

30. Poggetti V, Salerno S, Baglini E, Barresi E, Da Settimo F, Taliani S. Carbonic Anhydrase Activators for Neurodegeneration: An Overview. Molecules. 2022 Apr 14;27(8):2544. doi: 10.3390/molecules27082544. Accessed March 27, 2023.

31. Raghavan N, Miller AN, Gardner M, FitzGerald PC, Kerlavage AR, Johnston DA, Lewis FA, Knight M. Comparative gene analysis of Biomphalaria glabrata hemocytes pre- and post-exposure to miracidia of Schistosoma mansoni. Mol Biochem Parasitol. 2003 Feb;126(2):181–91. doi: 10.1016/s0166-6851(02)00272-4. Accessed April 8, 2023.

32. Rothwell PM, Wilson M, Elwin CE, Norrving B, Algra A, Warlow CP, Meade TW. Long-term effect of aspirin on colorectal cancer incidence and mortality: 20-year follow-up of five randomised trials. Lancet. 2010 Nov 20;376(9754):1741-50. doi: 10.1016/S0140-6736(10)61543-7. Accessed April 5, 2023.

33. Schistosomiasis. The World Health Organization. 2023. https://www.who.int/news-room/fact-sheets/detail/schistosomiasis. Accessed April 5, 2023.

34. Schmidt J, Oppermann E, Blaheta RA, Schreckenbach T, Lunger I, Rieger MA, Bechstein WO, Holzer K, Malkomes P. Carbonic-anhydrase IX expression is increased in thyroid cancer tissue and represents a potential therapeutic target to eradicate thyroid tumor-initiating cells. Mol Cell Endocrinol. 2021 Sep 15;535:111382. doi: 10.1016/j.mce.2021.111382. Accessed April 3, 2023.

35. Sievers F, Wilm A, Dineen D, et al. Fast, scalable generation of high-quality protein multiple sequence alignments using Clustal Omega. Molecular Systems Biology. 2011 Oct;7:539. DOI: 10.1038/msb.2011.75. Accessed April 13, 2023.

36. Smith M, Yadav S, Fagunloye OG, Pels NA, Horton DA, Alsultan N, Borns A, Cousin C, Dixon F, Mann VH, Lee C, Brindley PJ, El-Sayed NM, Bridger JM, Knight M. PIWI silencing mechanism involving the retrotransposon nimbus orchestrates resistance to infection with Schistosoma mansoni in the snail vector, Biomphalaria glabrata. PLoS Negl Trop Dis. 2021 Sep 8;15(9):e0009094. doi: 10.1371/journal.pntd.0009094. Accessed February 27, 2023.

37. Sostres C, Gargallo CJ, Lanas A. Aspirin, cyclooxygenase inhibition and colorectal cancer. World J Gastrointest Pharmacol Ther. 2014 Feb 6;5(1):40–9. doi: 10.4292/wjgpt.v5.i1.40. Accessed April 5, 2023.

38. Sudsarn P, Boonmars T, Ruangjirachuporn W, Namwat N, Loilome W, Sriraj P, Aukkanimart R, Nadchanan W, Jiraporn S. Combination of Praziquantel and Aspirin Minimizes Liver Pathology of Hamster Opisthorchis viverrini Infection Associated Cholangiocarcinoma. Pathol Oncol Res. 2016 Jan;22(1):57–65. doi: 10.1007/s12253-015-9967-y. Accessed April 14, 2023.

39. Supuran CT. Carbonic anhydrases: novel therapeutic applications for inhibitors and activators. Nature Reviews Drug Discovery 7, no. 2 (2008): 168–181. doi: 10.1038/nrd2467. Accessed April 13, 2023.

40. Svastová E, Hulíková A, Rafajová M, Zat’ovicová M, Gibadulinová A, Casini A, Cecchi A, Scozzafava A, Supuran CT, Pastorek J, Pastoreková S. Hypoxia activates the capacity of tumor-associated carbonic anhydrase IX to acidify extracellular pH. FEBS Lett. 2004 Nov 19;577(3):439–45. doi: 10.1016/j.febslet.2004.10.043. Accessed April 11, 2023.

41. Verjee MA. Schistosomiasis: Still a Cause of Significant Morbidity and Mortality. Res Rep Trop Med. 2019 Dec 31;10:153–163. doi: 10.2147/RRTM.S204345. Accessed March 8, 2023.

42. WHO. (2018). Schistosomiasis: Progress report 2001–2011, strategic plan 2012–2020. Geneva, Switzerland: World Health Organization. Retrieved from https://www.who.int/schistosomiasis/resources/9789241503174/en/. Accessed March 15, 2023.

43. Zolfaghari Emameh R, Barker HR, Turpeinen H, Parkkila S, Hytönen VP. A reverse vaccinology approach on transmembrane carbonic anhydrases from Plasmodium species as vaccine candidates for malaria prevention. Malar J. 2022 Jun 15;21(1):189. doi: 10.1186/s12936-022-04186-7. Accessed April 11, 2023.

